## Supplementary information for "Chloride intracellular channel (CLIC) proteins function as fusogens"

[1] Department of Physiology and Pharmacology, Faculty of Medicine, Tel-Aviv University, Tel-Aviv, 6997801, Israel

[2] School of Chemistry, Raymond & Beverly Sackler Faculty of Exact Sciences, Tel Aviv University, 6997801 Tel Aviv, Israel

[4] Department of Cell and Developmental Biology, Faculty of Medicine, Tel-Aviv University, Tel-Aviv, 6997801, Israel

[6] Tel Aviv Sourasky Medical Center, Tel Aviv, 6423906, Israel

[7] Sagol School of Neuroscience, Tel Aviv University, Tel Aviv, 6997801, Israel

[*] These authors contributed equally

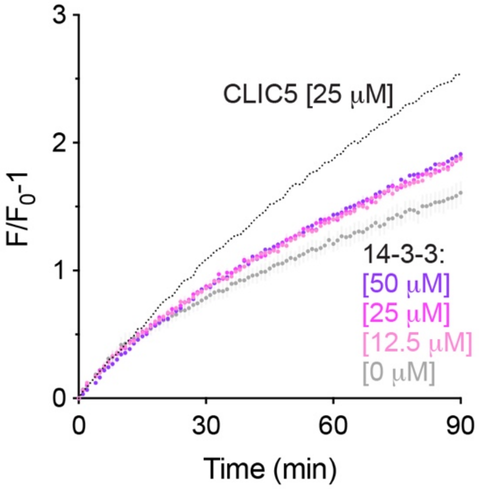


**Supplementary Figure 1. 14-3-3 does not induce lipid mixing.** Dose-response analysis of liposomal membrane mixing by 14-3-3 using the R18 fluorescence unquenching assay. CLIC5 (25 μM; Fig. 3a) is shown as a positive control.

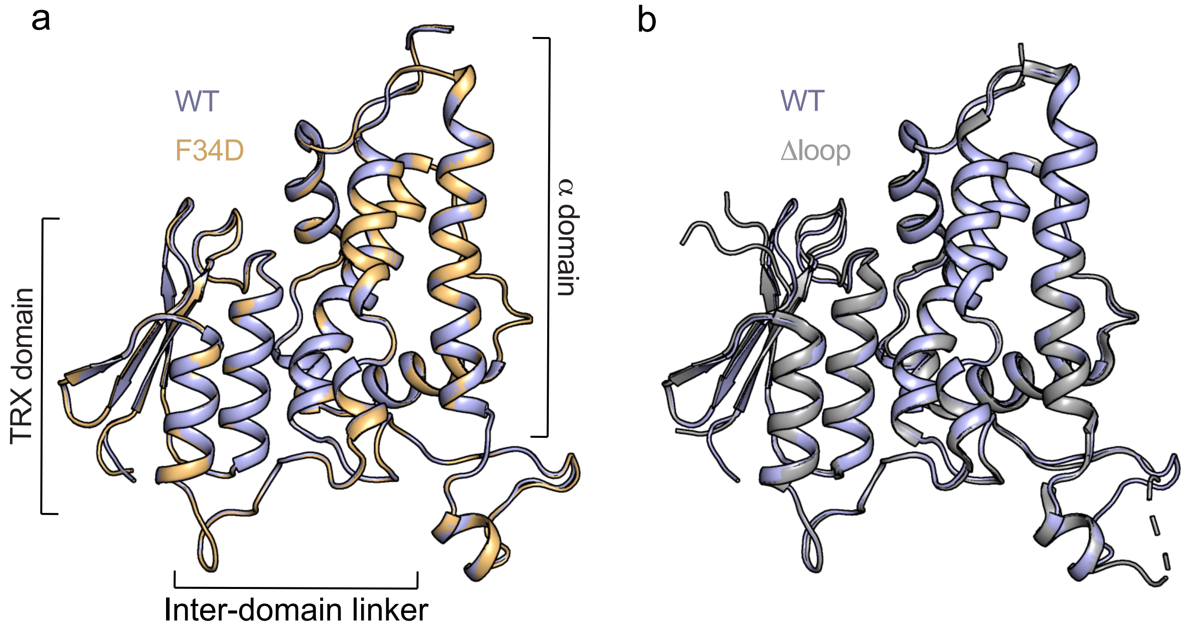


**Supplementary Figure 2. CLIC5-F34D does not induce significant structural perturbations.**

**(a, b)** Superposition of CLIC5-WT (PDB 6Y2H) and CLIC5-Δ57-68-F34D (a) or CLIC5-Δ57-68-WT (b). Neither the mutation or the deletion of the loop results in major structural alterations.

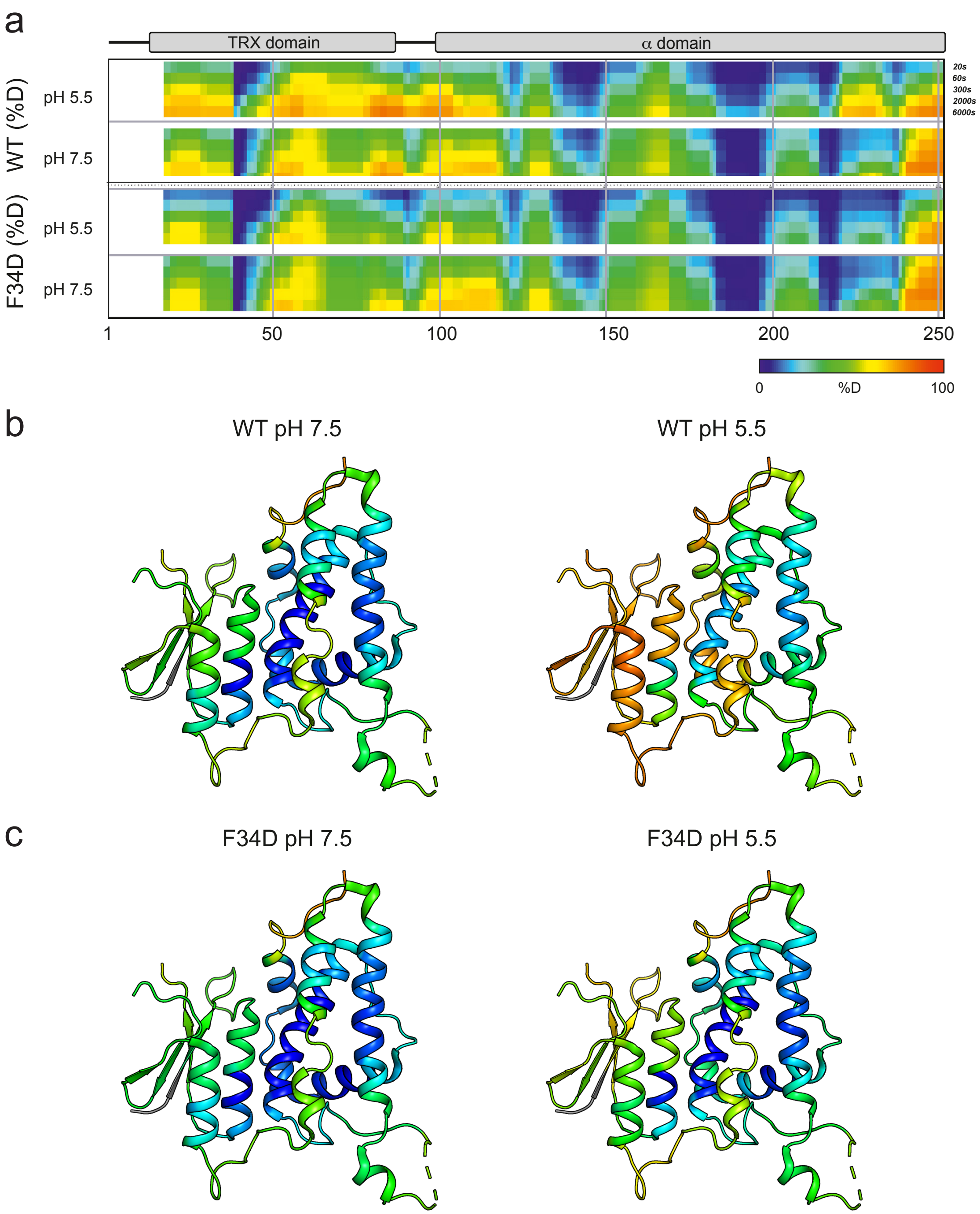


**Supplementary Figure 3. Deuterium uptake profiles of CLIC5-WT and CLIC5-F34D. (a)** Deuteration levels at the indicated time points for CLIC5-WT (upper panels) and CLIC5-F34D (lower panels), at the indicated pH values. **(b, c)** The deuterium uptake levels at 60 seconds (pH 7.5) or 6000 seconds (pH 5.5) are projected onto the structure of CLIC5-WT (PDB 6Y2H), as indicated.

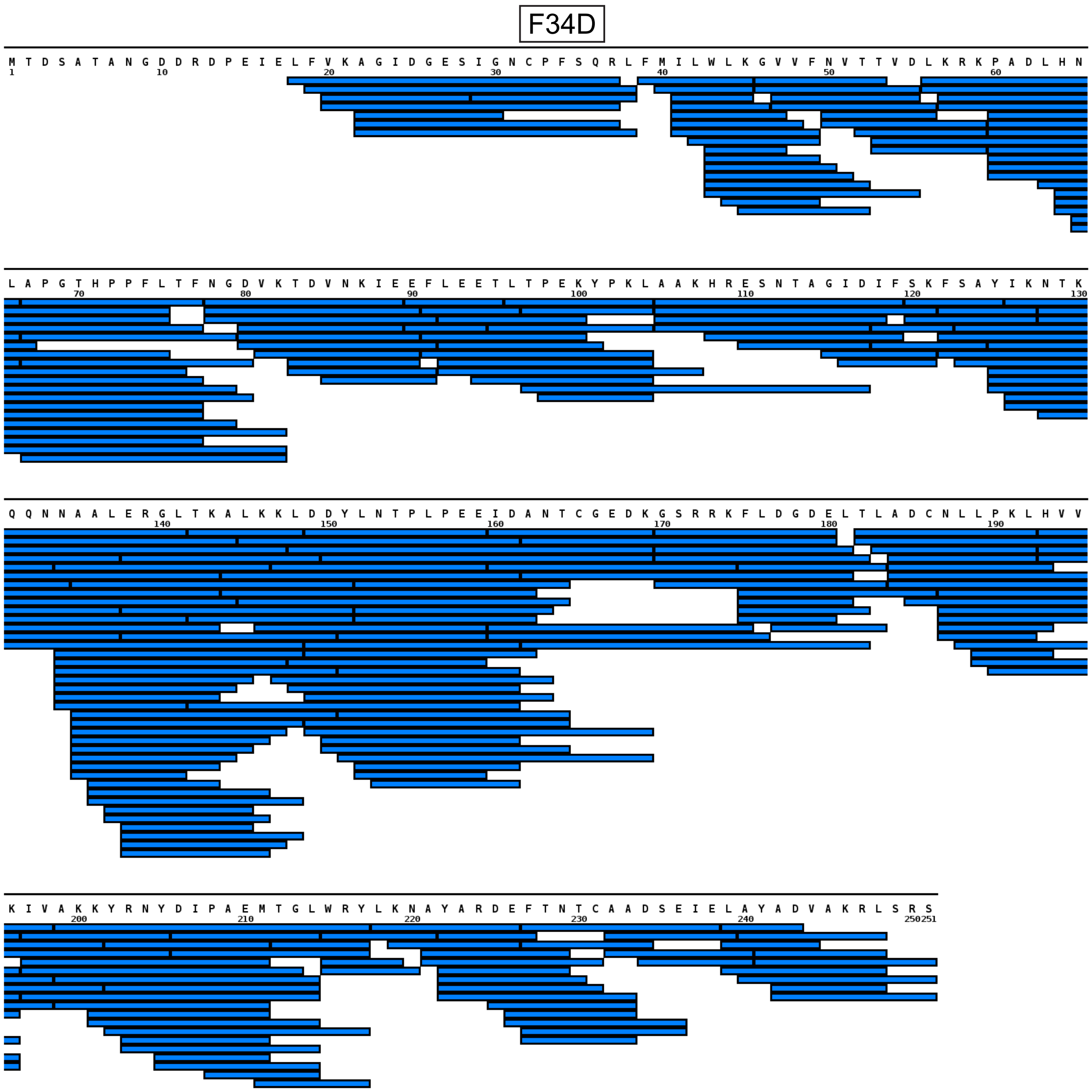


**Supplementary Figure 4. Sequence coverage in HDX-MS experiments.** The coverage map shows peptides generated after online digestion using immobilized pepsin/nepenthesin-2 under HDX-MS conditions.

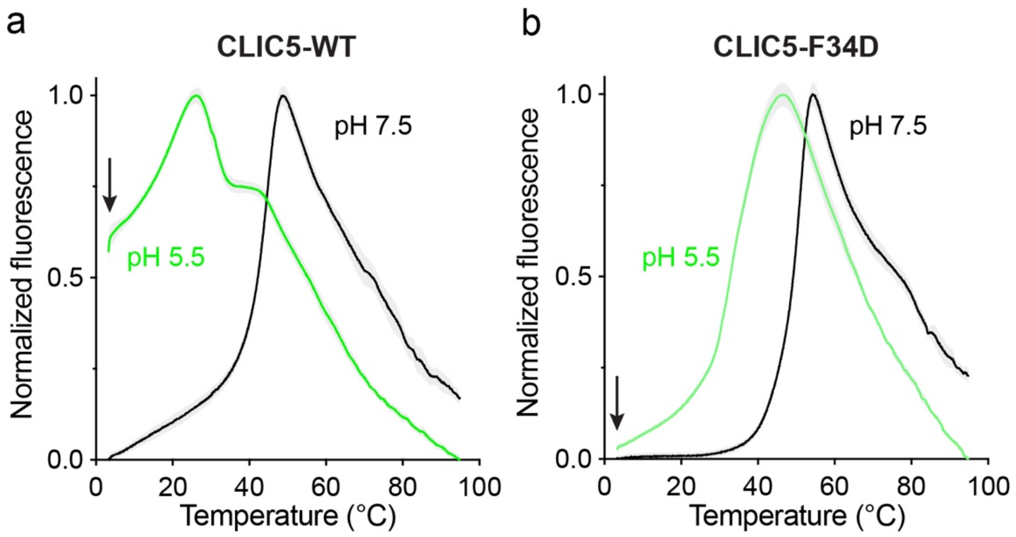


**Supplementary Figure 5. Thermal shift assay (TSA) analysis of CLIC5-WT and CLIC5-F34D. (a, b)** Normalized SYPRO Orange fluorescence-temperature relation of CLIC5-WT (a) and CLIC5-F34D (b) at the indicated pH values. Basal fluorescence signals are indicated (arrows).

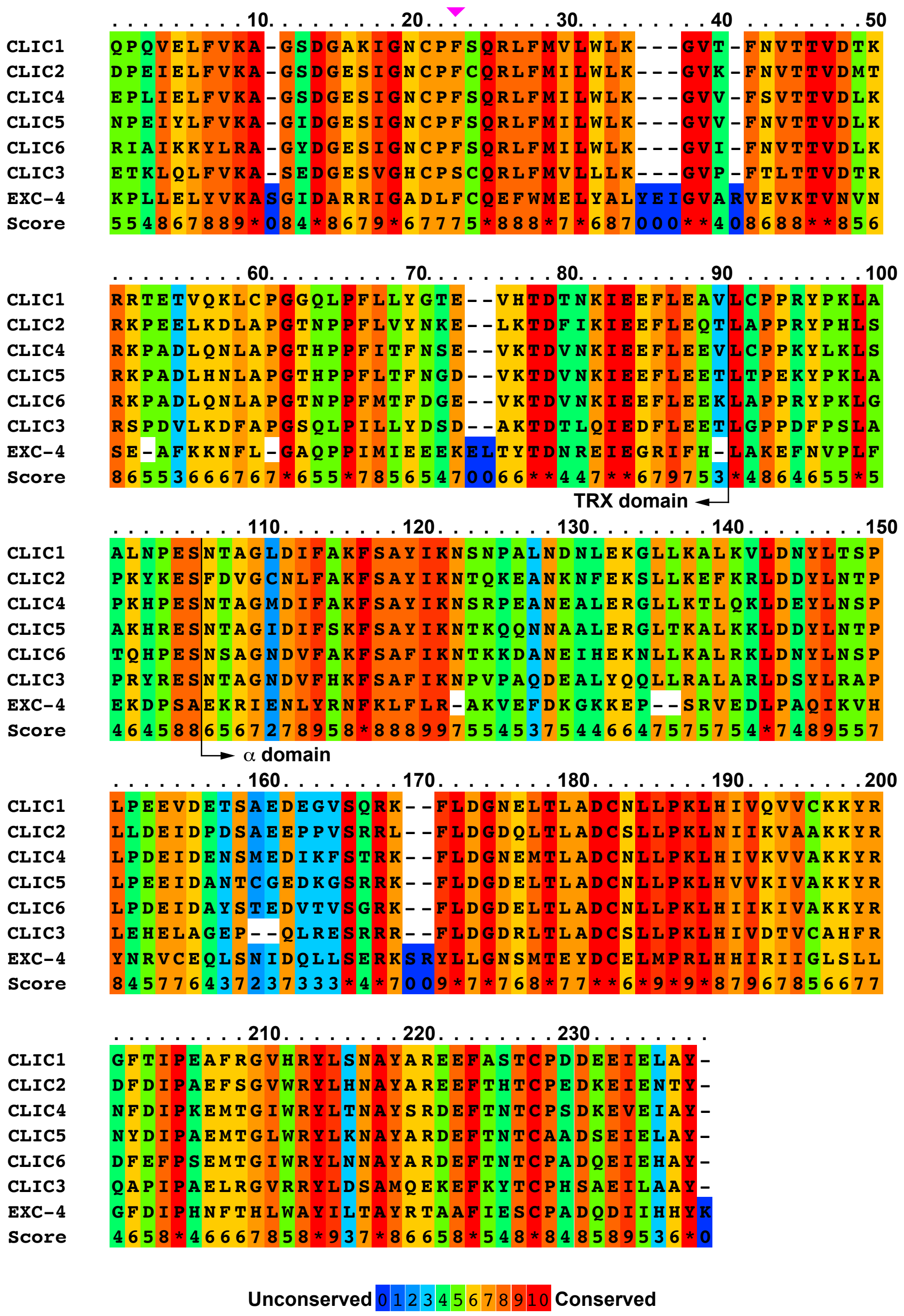


**Supplementary Figure 6. CLIC proteins sequence similarity.** CLIC domain PRALINE Multiple sequence alignment (https://www.ibi.vu.nl/programs/pralinewww/) of all human CLIC family members and EXC-4. TRX and α domain boundaries and CLIC5-F34 are indicated (black arrows and a purple arrowhead, respectively).
